# RNA origami scaffolds as a cryo-EM tool for investigating aptamer-ligand binding of a Broccoli-Pepper FRET pair

**DOI:** 10.1101/2022.08.25.505116

**Authors:** Néstor Sampedro Vallina, Ewan K.S. McRae, Bente Kring Hansen, Adrien Boussebayle, Ebbe Sloth Andersen

## Abstract

RNA nanotechnology uses motifs from nature as well as aptamers from in vitro selection to construct nanostructures and devices for applications in RNA medicine and synthetic biology. The RNA origami method allows cotranscriptional folding of large RNA scaffolds that can position functional motifs in a precise manner, which has been verified by Förster Resonance Energy Transfer (FRET) between fluorescent aptamers. Cryogenic electron microscopy (cryo-EM) is a promising method for characterizing the structure of larger RNA nanostructures. However, the structure of individual aptamers is difficult to solve by cryo-EM due to their low molecular weight. Here, we place aptamers on the RNA origami scaffolds to increase the contrast for cryo-EM and solve the structure of a new Broccoli-Pepper FRET pair. We identify different modes of ligand binding of the two aptamers and verify by selective probing. 3D variability analysis of the cryo-EM data show that the relative position between the two bound fluorophores on the origami fluctuate by only 3.5 Angstrom. Our results demonstrate the use of RNA origami scaffolds for characterizing small RNA motifs by cryo-EM and for positioning functional RNA motifs with high spatial precision. The Broccoli-Pepper apta-FRET pair has potential use for developing advanced sensors that are sensitive to small conformational changes.

## INTRODUCTION

The structural versatility of RNA makes it an ideal substrate for the design of functional nanodevices for applications in biotechnology and medicine. RNA’s structural properties and ability to interact with proteins, other nucleic acids, and small molecules, allows for the precise spatial scaffolding of different molecular components ^1-3^. Functional RNA scaffolds have been developed by incorporating ribozymes ^4^, riboswitches ^5^ and small-molecule binding aptamers ^6,7^. RNA scaffolds bearing protein-binding domains have the potential to be genetically expressed and serve for synthetic biology purposes by localizing enzymes together to increase metabolic flux ^8,9^ or gene expression regulation ^10-12^. RNA origami is a single-stranded RNA architecture based on the folding of RNA during transcription ^13,14^. Produced isothermally, RNA origami nanostructures can be folded in high yield and be used for constructing genetically encodable RNA nanodevices. The modular design principles of this architecture allow for the precise spatial arrangement of different RNA motifs, which allows for the development of functional scaffolds ^6,11,12^.

Using Cryo-EM, many biological structures have been solved with near atomic level resolution ^15-17^. Large macromolecular complexes are more prone to favourable data acquisition due to high signal-to-noise ratio and identifiable asymmetric features. Smaller complexes (<50 kDa) remain difficult to visualize by cryo-EM due to low contrast and high background. RNA structures as small as 28 kDa have been recently solved using cryo-EM thanks to optimized sample preparation protocols ^18,19^. A strategy based on homo-oligomerization of the target using kissing loops has recently been developed to increase molecular weight and mitigate flexibility ^20^. Following this method, the structure of RNA targets such as the Tetrahymena group I Intron, the Azoarcus group I intron and the FMN riboswitch were solved at high resolution. An alternative strategy that has been used for proteins is to scaffold smaller structural targets on larger structures ^21,22^. This can help optimize the folding and homogeneity of low molecular weight targets, as well as increase the signal-to-noise ratio while not interfering with the flexibility of the molecule. This strategy may also serve as a method for solving small RNA targets.

Förster resonance energy transfer (FRET) represents a versatile tool to characterize scaffolding effects and build devices sensitive to small structural changes, having applications in studies of localization of molecules and conformational changes ^23-26^. Fluorogenic aptamers (FAs) are RNA sequences that interact with and activate fluorophores, reducing their non-radiative decay through vibrational and rotational relaxation by rigidifying their aromatic orbital systems ^27,28^. RNA devices that incorporate FAs can be used to investigate structural dynamics of RNAs and to develop biosensors. We previously functionalized an RNA origami tile with Spinach and Mango aptamers, whose complexes with DFHBI and YO3-biotin acted as a FRET donor and acceptor, respectively (named an apta-FRET system) ^6^. The RNA origami scaffold with FAs rigidly incorporate the fluorophores leading to both a distance- and angle-dependency of the FRET efficiency ^29^. Apta-FRET has also been demonstrated with the iSpinach and Mango-IV aptamers joined together by a single-stranded region ^30^. Jeng et al. observed angular changes in an apta-FRET system and were able to sense the structural stabilization of a metal-ion binding riboswitch ^26^. Geary et al. used RNA origami to demonstrate controlled positioning of Spinach and Mango aptamers and effects of helix length and flexibility on FRET efficiency ^14^. All of the above apta-FRET systems used YO3-biotin as acceptor, however, this fluorophore has limited stability and high background when internalized by cells ^6^ and its fluorescence has been shown to be weakly activated by Spinach ^30^. Therefore, new aptamers and fluorophores with stable and bright properties are needed as acceptors to develop RNA FRET devices that can act inside cells.

Here, we use RNA origami to construct a new FRET pair using FAs, where the Broccoli/DFHBI-1T complex acts as a donor and the Pepper/HBC620 complex acts as an acceptor, and by tuning their relative positioning we obtain high FRET. Using cryo-EM single particle averaging methods, we reconstruct the apo and fluorophore-bound states of our RNA origami FRET pair to 4.5 Angstrom resolution. Supported by SHAPE probing experiments, we find that the Broccoli aptamer does not change shape upon ligand binding, while Pepper is rigidified upon ligand binding. Finally, we use 3D variability analysis of the particles isolated from the cryo-EM data to model the positional variance of the two fluorophores in the RNA scaffold and find that the Förster radius varies by only 3.5 Angstrom and is dominated by translational / non-rotational modes of movement. Our results demonstrate the use of a new FRET pair with FAs more suitable for in vivo applications and show that RNA origami scaffolds can be used for characterizing small RNA motifs by cryo-EM and for positioning functional RNA motifs with high spatial precision.

## RESULTS

### FRET between Broccoli and Pepper aptamers

To develop a new apta-FRET system with improved properties we suggest using the Broccoli and Pepper aptamers. The Broccoli aptamer is a shorter version of Spinach with improved folding in vivo ^31^ which, in complex with the DFHBI-1T fluorophore, has a reported excitation maximum at 485 nm and emission maximum at 505 nm ^32^. The Pepper aptamer in complex with the fluorophore HBC620 has shown excellent fluorescent properties in vitro and in vivo, with reported excitation and emission maxima at 577 nm and 620 nm, respectively ^33^. When placed in proximity, these two aptamers in complex with their cognate fluorophores represent a promising FRET pair candidate (Fig. 1A), while being also potentially expressible and stable inside the cellular milieu.

**Fig. 1.**
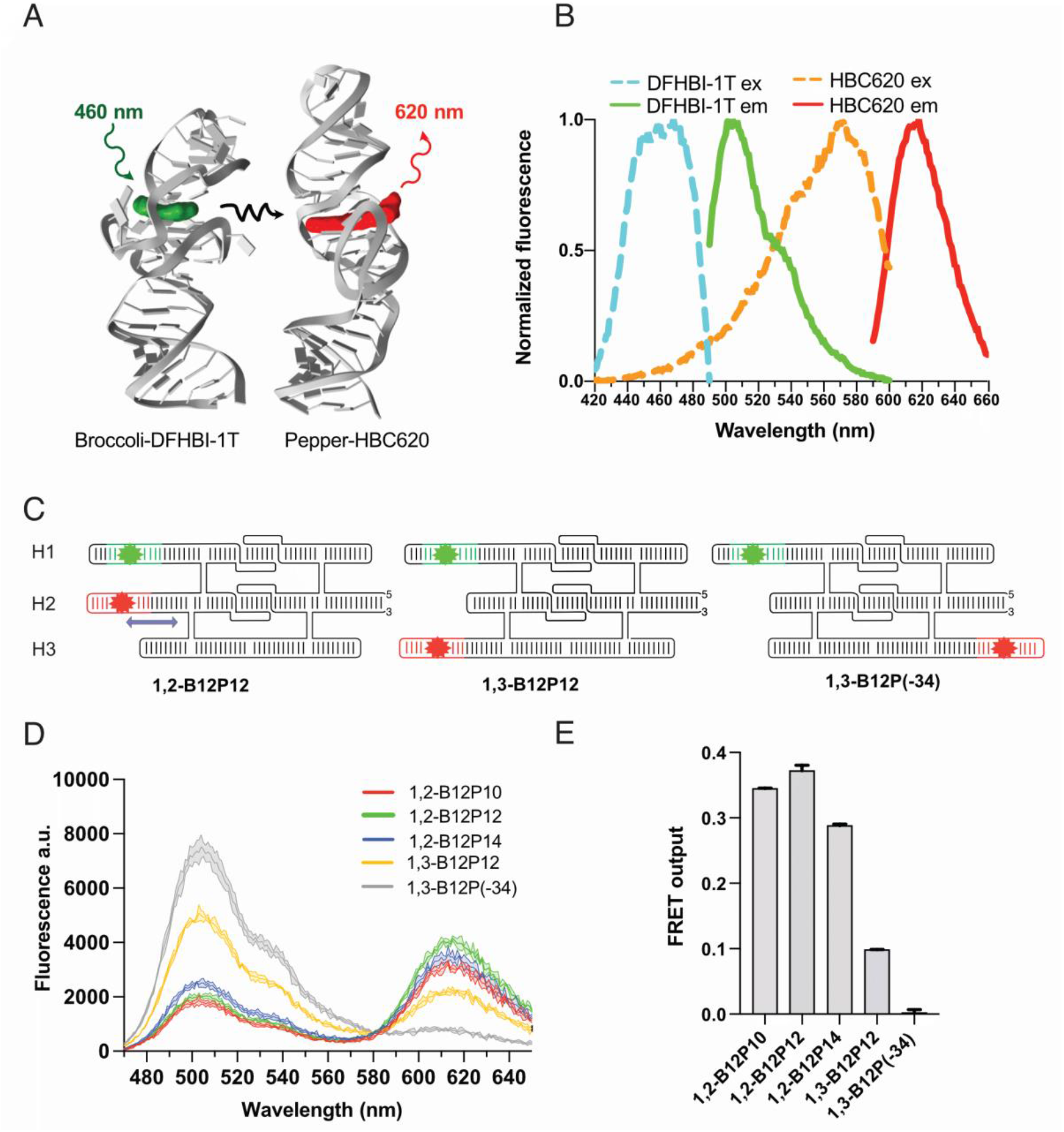
FRET between Broccoli and Pepper aptamers. (A) Structural model of Broccoli and Pepper aptamers shown in cartoon format with their cognate fluorophores DFHBI-1T (green) and HBC620 (red) shown as spheres. Excitation, energy transfer and emission illustrated as wavy lines. (B) Measured excitation and emission spectra of DFHBI-1T and HBC620 in complex with their cognate aptamers. (C) Depiction of RNA origami tiles with different arrangements of the fluorogenic aptamers. (D) FRET output measured after 30 mins upon addition of the fluorophores (1 µM) to the RNA origami tiles (100 nM). Measured fluorescence spectra at 460 nm excitation. Data corresponds to 3 technical replicates, shown as mean ± SD. (E) Calculated absolute FRET output measured at 460 nm excitation and 620 nm emission. Data corresponds to 3 technical replicates, shown as mean ± SD.

We first experimentally analyzed the spectral properties of DFHBI-1T bound by Broccoli and HBC620 bound by Pepper (Fig. 1B). DFHBI-1T/Broccoli emission spectra and HBC620/Pepper excitation spectra were found to have a significant overlap, which is beneficial for FRET to occur. Furthermore, the broad excitation spectra of DFHBI-1T/Broccoli allows for excitation of the donor at 460 nm with negligible excitation of the acceptor (HBC620/Pepper). The minimal overlapping excitation spectra between DFHBI-1T and HBC620 results in minimal direct excitation of the acceptor (Fig. 1B, Supplementary Table 2). To obtain FRET, we incorporated the Broccoli and Pepper aptamers onto a 3-helix

RNA origami scaffold (Fig. 1C), similarly to the 2-helix apta-FRET scaffolds used by Jepsen et al. ^6^. We designed five scaffolds with different aptamer placements to investigate the effect of donor and acceptor aptamer distance on FRET efficiency. The apta-FRET constructs are annotated as x,y-Bz-Pw, where x and y refer to the helix segment on which an aptamer is placed, and z and w refers to the distance in base pairs from the crossover on the RNA origami scaffold to Broccoli (B) and Pepper (P), respectively. In three constructs (1,2-B12P10, 1,2-B12P12, 1,2-B12P14), the aptamers were placed on helices 1 and 2 to put the fluorophores in close proximity, while varying the length of the stem before the Pepper aptamer (arrow in Fig. 1C) and keeping the Broccoli stem at 12 bp. In two constructs, the aptamers were placed on helices 1 and 3; from our in silico 3D modelling, 1,3-B12P12 places the aptamers at ∼2 nm distance while 1,3-B12P(−34) places the aptamers at ∼16 nm distance, the latter being outside FRET distance. The secondary structures and sequences for each design can be found in Supplementary Table 1.

The highest FRET output of 37 ± 0.4 % was observed for the 1,2-B12-P12 design, since moving the Pepper aptamer to either the left (1,2-B12P10) or right (1,2-B12P14) of this position resulted in reduced FRET efficiency of 34 ± 0.04 % and 29 ± 0.1 %, respectively (Fig. 1 D,E, Supplementary Table 2). This decrease can either result from a distance effect or an oriented dipole effect as has been documented previously ^6,26^. When placing the aptamers further apart on helix 1 and 3 in the 1,3-B12P12 design, we observe a FRET output of ∼10% and when placing the aptamers outside FRET distance in the 1,3-B12P(−34) design, no FRET was observed (Fig. 1 D,E, Supplementary Table 2). In conclusion, we have shown that DFHBI-1T/Broccoli and HBC620/Pepper function as a FRET pair with high FRET efficiency when located at an appropriate distance and orientation.

### Cryo-EM structure of Broccoli-Pepper scaffold in apo and bound states

To better understand the relative positioning of the fluorescent aptamers on our RNA origami scaffold, we used cryo-EM single particle averaging methods to determine the structure of the 1,2-B12P12 scaffold with and without fluorophores bound. Our cryo-EM data set for the Bound scaffold contained 5354 movies, resulting in a refined particle stack containing 150,704 particles that produced a reconstruction with a GSFSC (0.143) resolution estimate of 4.43 Angstrom (Supplementary Fig. 1 and Supplementary Table 3). The Apo scaffold dataset had only 1605 movies and resulted in a refined particle stack with 50,868 particles reaching a GSFSC (0.143) of 4.55 Angstrom (Supplementary Fig. 2).

**Fig. 2.**
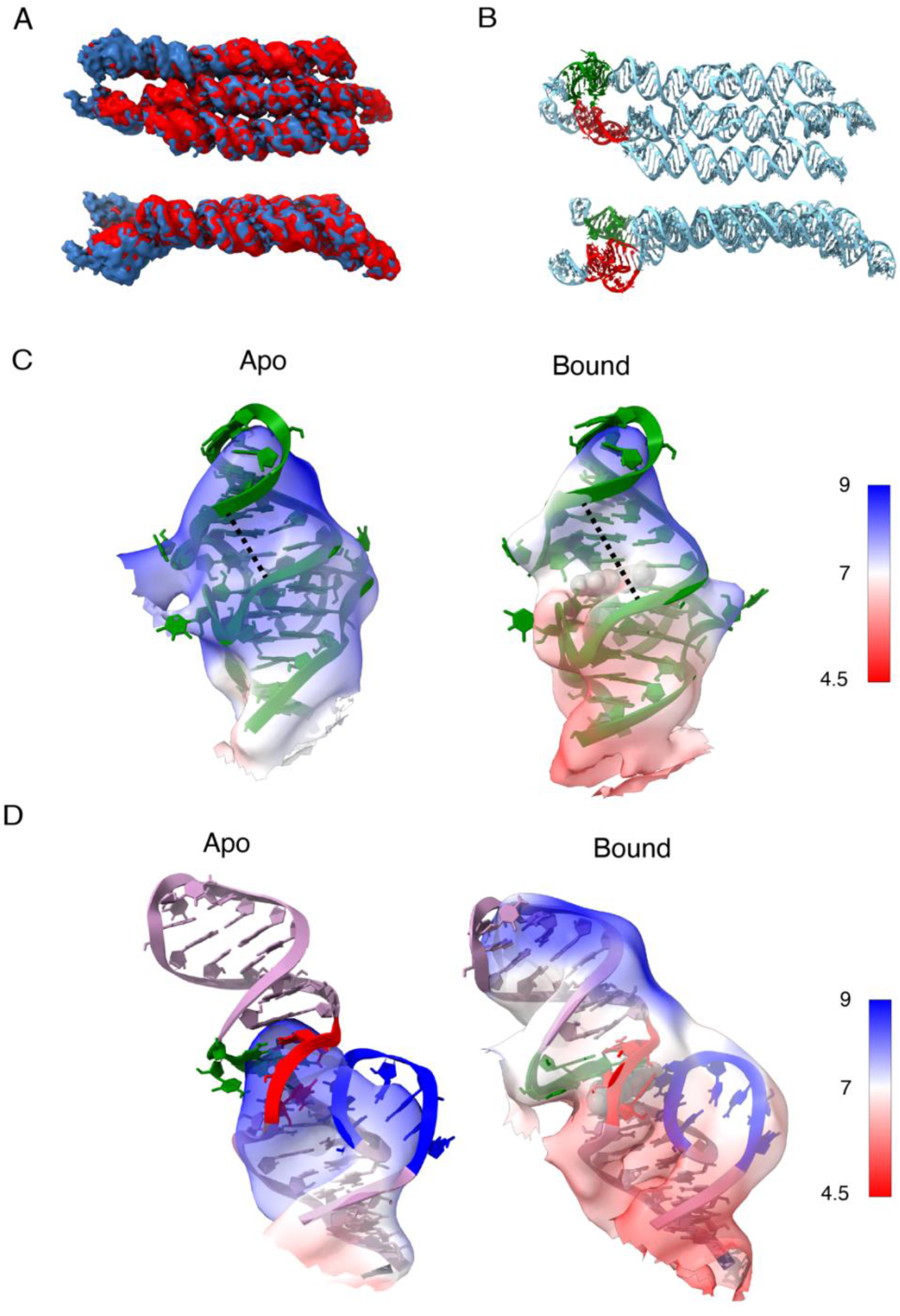
Cryo-EM structure of Pepper and Broccoli aptamers in apo and bound states. (A) Overlay of the cryo-EM maps of the apo (red) and ligand bound (blue) 1,2-B12P12. (B) Atomistic model built into the ligand bound cryo-EM map of 1,2-B12P12 showing the Brocolli (green) and Pepper (red) aptamer locations. (C) Close up view of the Brocolli aptamer in the apo and DFHBI bound cryo-EM maps. (D) Close up view of the Pepper aptamer in the apo and HBC620 bound cryo-EM maps.

The Apo- and Bound-scaffold overlap very well, with only slight positional variability in the aptamer regions (Fig. 2A). An unexpected curvature in the central helix of the scaffold results in the Pepper aptamer being below the plane of the scaffold (Fig. 2B, bottom, red motif). A slight curvature is also introduced at the stem of the Broccoli aptamer positioning it slight above the plane of the scaffold (Fig. 2B, bottom, green motif). Although the local resolution of the Broccoli aptamer ranges from 8.0 to 9.5 Angstrom for the apo state and 5.5 to 8.7 for the bound state, the overall shapes of the two states are similar, indicating that the G-quadruplex structure is not affected by the ligand binding (Fig. 2C). Removal of the ligand from our Broccoli model and refinement into our Apo-map results in the displacement of the adenine-uracil base pair that normally stacks on top of the ligand. This base pair now stacks directly on top of the final G-quartet, changing the angle of the terminal helix and, consequently, the major groove opposing the ligand-binding pocket is narrowed from 13.6 Angstrom in the Bound model to 9.2 Angstrom in our Apo model (indicated by dashed line in Fig. 2C).

In contrast, the Pepper aptamer is missing or has weak signal from key ligand-binding regions in the apo state that are clearly present in the bound state (Fig. 2D). Specifically, J3/2 (Fig. 3A), which forms the side of the ligand binding pocket, has weaker signal than the bound state at similar map thresholds. The local resolution for J3/2 reaches 7 Angstrom for the bound state, but only 9 Angstrom for the apo state. Furthermore, at a map threshold approximating 9 Angstrom resolution, 145 atoms from J3/2 are outside the contour of the Apo map. Whereas at a map threshold approximating 7.5 Angstrom, 0 atoms from J3/2 are outside the contour of the Bound map. At this threshold level of ∼9 Angstroms the Apo reconstruction terminates at the ligand binding site and the P1 helix is not observed until the threshold is extended to observe features of up to 10.5 angstrom local resolution. In comparison, the entire P1 helix is contained within the bound state reconstruction at a threshold level equivalent to 8.5 Angstrom local resolution. In conclusion, the cryo-EM data allows us to observe a major rigidification of the Pepper aptamer upon ligand binding.

**Fig. 3.**
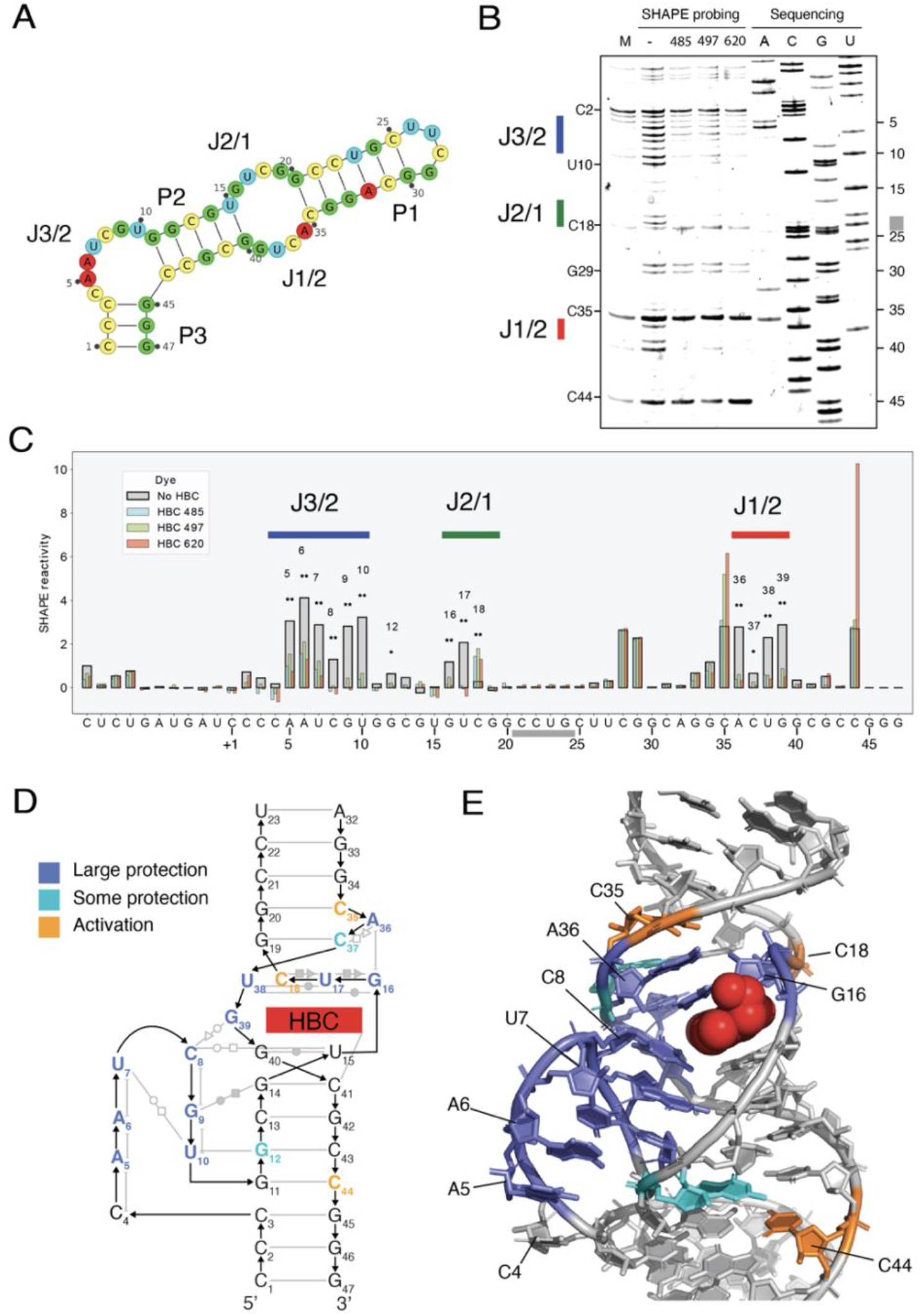
SHAPE probing of the Pepper aptamer in apo and bound states. (A) Secondary structure blueprint for Pepper with the labelling used in the text. (B) SHAPE gel analysis of pepper aptamer in the apo and HBC 485, 497 and 620 bound states. Grey marking in sequencing lane indicates compressed area. (C) Quantitative per-nucleotide SHAPE reactivity analysis for Pepper aptamer in the apo and ligand bound states. Signals are normalized by the signal at the non-binding C28 position. (D) Tertiary structure of the Pepper aptamer (PDB ID: 7EOP). Structure diagram showing tertiary elements: Base pairs are shown as grey lines with Leontis-Westhof annotation of non-Watson-Crick base pairs. Base pair planes are indicated by horizontal alignment. Stacking is indicated by vertical alignment. Protection is marked as colors on nucleotides. (E) Atomic structure shown with protection colored on nucleotides showing that the whole binding pocket gets stabilized upon ligand binding. HBC 620 shown in red sphere representation.

Comparing the per-residue cross-correlation (CC) values from our models and their respective EM reconstructions shows that we can model each residue with similar confidence in both the apo and bound state, except for the Pepper aptamer, which has lower CC values in the Apo model (Supplementary Fig. 3). Although these differences observed in the Pepper aptamer from our cryo-EM analysis could be due to a lesser number of particles being available during data processing for the apo state, the similarity in the Broccoli-containing regions and subsequent SHAPE analysis confirms what we have observed in the EM reconstructions.

### SHAPE probing of Pepper reveals cooperative binding of HBC

We performed SHAPE analysis on the Pepper aptamer in the absence and presence of 3 Pepper ligands: HBC485, HBC497, and HBC620 (Fig. 3 A,B, Supplementary Fig. 6). The junction regions J3/2, J2/1 and J1/2 were observed to have high SHAPE reactivity in the absence of the ligand and low SHAPE reactivity in the presence of the ligands indicating that the junction nucleotides cooperatively bind the ligands (Fig. 3 B,C). Inside the junction regions, we observed negligible difference in SHAPE reactivity between the three ligands tested, confirming that the mode of binding is conserved as indicated by the previously determined crystal structures ^34^.

Low SHAPE reactivity was observed for the first nucleotide (C4) from J3/2 both in the absence and presence of ligands (Fig. 3 B,C). From the crystal structure ^34^ and our EM data we would expect this to be one of the more dynamic residues as it has no hydrogen bonding partner and is only supported by base stacking from one adjacent nucleotide. The three next nucleotides (position 5-7) show significant flexibility in the apo-structure that is attenuated in the ligand bound state. The final three nucleotides of J3/2 (position 8-10) show complete loss of SHAPE reactivity upon ligand binding, supporting their interaction in the deep groove of P2 and role in forming the ligand binding site. C8 shows the lowest reactivity of these nucleotides in the apo state, indicating that it could be transiently sampling the ligand bound conformation.

The U15-G40 base pair forms the bottom of the ligand binding site and is stable in both the apo and bound states. C8 and G39 form the left side of the binding pocket (Fig. 3D,E) and are both SHAPE reactive in the apo state but less reactive in the presence of ligand (Fig. 3C). The top of the ligand binding pocket is formed by the mixed-base tetrad G16-U17-C18-U38 (Fig. 3D,E). G16 and U17 from J2/1 as well as U38 from J1/2 are SHAPE reactive in the unbound state but lose reactivity in the bound state, while C18 from J2/1 is unreactive in the apo state but reactive in the bound state (Fig. 3C). This suggests that C18 is stacked inside the helix in the absence of ligand but gets displaced when the ligand enters its binding site. In the crystal structure, C18 has one hydrogen bond to U17, but is on the exterior of the helix with no stacking partner (Fig. 3E) and is flexible as indicated by a high B-factor. The mixed-base tetrad is stabilized by stacking on the G19-C37 base pair. These nucleotides are both unreactive in the unbound and bound state, indicating that the base pairing observed in the crystal structure is maintained in the absence of ligand. A36 from J1/2 is reactive in the apo form and stabilized in the bound state, confirming its role in stabilizing the binding pocket by stacking on top of G16 (Fig. 3D,E).

Nucleotide C35 and C44 have an apparent increased SHAPE reactivity in the presence of some of the ligands, but since these positions are involved in base pairs in the crystal structure, they should not be SHAPE reactive. We observe bands corresponding to C35 and C44 in the mock lane (M in Fig. 3B) that are likely the result of premature termination of the reverse transcriptase (RT) at stable RNA structures. This is supported by C44 being positioned at the 3’ of P2 and C35 at the 3’of P1. The C35 and C44 bands appear more intensely in the Benzoyl Cyanide treated samples, suggesting an increase in termination due to the chemical modification. For C44, it is observed that HBC485 and HBC487 have similar intensity as with no ligand, but that HBC620 has a higher signal, which can be explained by its stronger binding affinity (HBC485 KD=8.0 nM, HBC497 KD=6.7 nM, HBC620 KD=6.1 nM) ^34^. For C35, we observe that HBC485 terminates at a similar level to no ligand, while HBC487 and HBC620 have a higher signal. Again, this fits with the binding strengths of the fluorophores.

We found that the Pepper junction regions are flexible in the apo state and become more structured in the bound state. The bound state fits very well the crystal structure. Also, from RT termination we see evidence of differential stabilization by the ligands. Our SHAPE data shows that even though the P1 helix is not apparent in our apo cryo-EM map, it is stable in the apo state. The lack of rigidity from the J3/2, J2/1 and J1/2 nucleotides likely result in a large amount of dynamics in the apo state that is averaged out during the single particle averaging analysis. Together, the cryo-EM and SHAPE data show that the Pepper aptamer undergoes significant structural rigidification upon ligand binding.

### Cryo-EM reveals conformational variability of FRET pair

In previous apta-FRET experiments it was observed that using longer stems to position the aptamers resulted on lower FRET efficiency, which suggests that the most variable regions of RNA origami are the termini of the helical components ^6,14^, bringing into question how precisely we can position the RNA motifs that we place at these variable positions. A partially cleaned particle stack from the Bound data set, with 241,297 particles, was used to perform 3D variability analysis (3DVA). Principal component analysis revealed three major movements, which corresponded to a density increase in the Pepper aptamer (PC0), movement of the Pepper aptamer (PC1) and movement of the Broccoli aptamer (PC2) (Fig. 4A, left, Supplementary Movie 1). The fluctuation of Pepper density observed in PC0 may correspond to the on-off binding of HBC620. For PC1 and PC2, we observe a continuous out-of-plane movement of the Pepper and Broccoli aptamers, respectively (Fig. 4A, right, Supplementary Movie 1). With the variability analysis we can identify the positional extrema of both the Pepper and Broccoli aptamers (Fig. 4B), which in turn allows us to determine the position of the fluorophores.

**Fig. 4.**
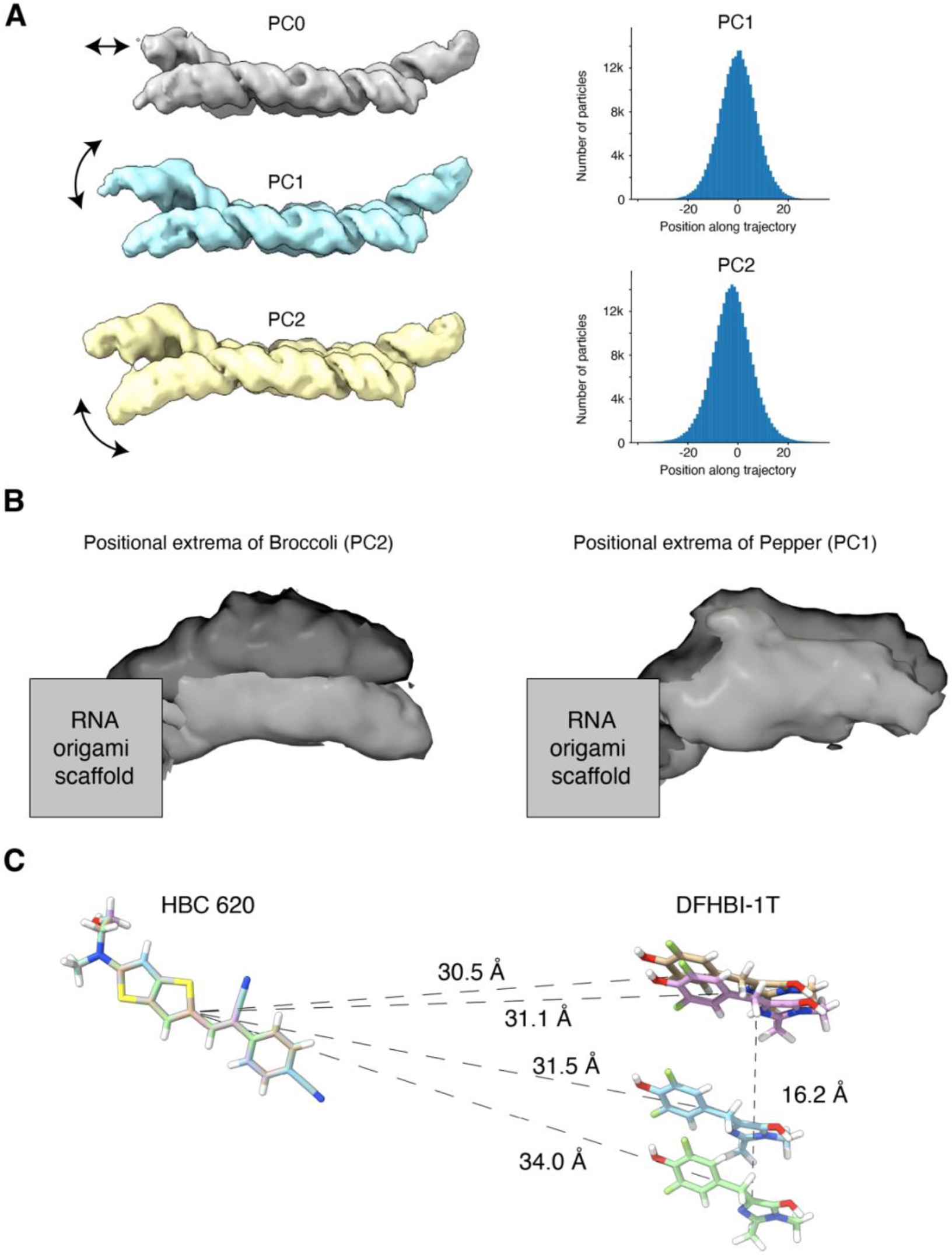
Cryo-EM 3D variability analysis of Pepper and Broccoli aptamers. (A) Representative structures are shown for the principal component analysis (PC0, PC1 and PC2). Arrows indicate the most prominent movements. Gaussian distribution of particles along two principal reaction coordinates (PC1, PC2) determined by 3DVA (right). (B) Intermediate reconstructions using particle subsets from the extremes of PC0 and PC1 show the positional variability of broccoli and pepper aptamers. (C) HBC ligands from pepper were aligned while maintaining the spatial relationship with the DFHBI ligand from Broccoli from the extrema reconstructions. Distances from HBC to the center of each DFHBI were measured as well as the furthest distance between DFHBI fluorophores.

By rigid-body fitting the crystal structures of the aptamers into the reconstructions we can determine the relative positions of the fluorophores for each of the extrema conformations. Then, while keeping the relative position of DFHBI-1T to HBC620 intact, we align the HBC620 models (Fig. 4C). For apta-FRET systems the fluorophore’s dipole moments are oriented in relation to each other and this orientation has a strong effect on FRET ^26,29^. We observe that the rotational orientation of the two fluorophores has little variation between the extrema positions, which is explained by the fixation of the aptamers on the RNA origami scaffold (Fig. 4B). The distance between fluorophores also has a strong effect on FRET efficiency ^29^.From our data we can see that the range of motion of a given fluorophore is close to ∼16 Angstrom. However, if we measure the distance between HBC620 and DFHBI-1T over the range of motion we see that the actual fluorophore distance ranges from 30.5 to 34.0 Angstrom. Thus, the range of probable distances for the two fluorophores has a variance of only ∼3.5 Angstrom, demonstrating that the current RNA origami paradigm allows us to position these two small molecules with sub-nanometer precision. Furthermore, since the distribution of particles across the reaction coordinates can be approximated as Gaussian (Fig. 4A, right), and therefore inform on the energy landscape of the particles, the majority of the population of molecules will be in an intermediate state, only sampling these extrema transiently.

## DISCUSSION

In this study we used RNA origami design tools ^14^ to position the Broccoli ^31^ and Pepper ^33^ aptamers at a close distance, generating a novel apta-FRET pair with comparable efficiency to our previously reported apta-FRET system ^6,14^. RNA origami design allows us to control the spatial orientation of the fluorophores and can thus be used to study the effects of dipole moment orientation and distance on FRET efficiency ^29^. The RNA origami designs used in this study arranges helices in a near-parallel manner, which results in a parallel positioning of aptamers and bound fluorophores. In another study, a metal-binding junction was used to place the aptamers at a non-parallel angle ^26^. By rationally designing RNA nanostructures of different geometries, it will be possible to explore the full range of dipole orientations. Apta-FRET systems can be used to create RNA devices that detect small conformational changes and be used to develop ratiometric biosensors ^35^. The apta-FRET system that we introduce in this study can also serve as basis for designing intracellular sensors, since the Broccoli and Pepper aptamers have both been independently verified to activate the fluorescence of their cognate fluorophores inside cells ^31,33,36^.

By combining cryo-EM and chemical probing methods we have obtained novel structural information about the unbound states of the Broccoli and Pepper fluorescent aptamers. This information can be further used to design improved Pepper aptamers with less flexibility, or to design a switchable Pepper aptamer where the flexible regions are sequestered by tertiary motifs, trapping the aptamer in an inactive state. Furthermore, the structural characterization of our RNA origami scaffolded apta-FRET system confirms that our in-silico design process can accommodate fluorescent aptamer motifs and still produce high-fidelity designer sequences that can fold cotranscriptionally into the predicted structure.

Due to low signal-to-noise ratios, small structural RNA targets can present a challenge for cryo-EM structure determination ^18^. Here, the RNA origami scaffold increases the size of the structural RNA targets and aids in the structural determination of the Broccoli and Pepper aptamers. Similarly, RNA origami can aid in structural determination of other interesting RNA motifs. An oligomerization approach using kissing loops has been used to characterize RNA-only targets by increasing their molecular weight and mitigate flexibility to aid in cryo-EM structure determination, obtaining a high resolution ^20^. However, similarly to crystallography, this approach can limit the natural flexibility and structural variability of the particles and even constrain them into artefactual conformations. In our approach, the RNA particles can be in their unconstrained solution conformation and therefore, the flexibility of the scaffold can be studied.

Since we obtained the cryo-EM map before the publication of the Pepper crystal structures ^34^, we attempted structure determination using the DRRAFTER pipeline ^37,38^. The models generated reached a mean pairwise RMSD (convergence) of 3.5 Angstrom, and out of the top 10 models, 9 had the correct strand path for J3/2. Since DRRAFTER cannot consider small molecule ligands, it was unable to recapitulate the ligand binding pocket and accurately place the ligand. When using a map simulated from the atoms at 5 Angstrom resolution in ChimeraX, the x-ray structure and DRRAFTER model both correlate comparably well (0.88) to our experimental EM map (data not shown). This shows that while de novo model building methods can accurately trace the backbone of complex RNA motifs into low resolution maps, caution should be taken when interpreting the base positions from such methods, especially in cases where non nucleic acid ligands are present.

In summary, the RNA origami architecture represents a versatile tool for scaffolding and combining RNA aptamers and other motifs with sub-nanometer precision, aiding in cryo-EM studies and presenting opportunities for further development of functional scaffolds.

## MATERIALS AND METHODS

### RNA sequence design

The RNA origami sequence design pipeline is extensively explained in Geary et al.^14^. Briefly, using a standard text editor, the different structural motifs were incorporated and routed on a single strand. The fluorogenic aptamers, as well as specific 3’ and 5’ -end primer binding regions ending in GGA (an optimal initiation sequence for the T7 RNA polymerase) were incorporated as sequence constrains. The sequences matching the specified constrains were then generated using the perl script “batch-revolvr.pl” from the ROAD package ^14^, available at https://github.com/esa-lab/ROAD.

### Synthesis of DNA templates

The DNA templates for the different RNA designs were produced by PCR amplification using Phusion High-Fidelity DNA polymerase (NEB) of double stranded gene fragments (gBlocks) synthetized by Integrated DNA Technologies (IDT). Amplifications were performed in 100 µl reactions containing 1X Phusion HF buffer (NEB), 1 µM of each primer (ordered from IDT), 200 µM dNTPs (Invitrogen), 4 ng of gBlock template and 1 Unit of Phusion DNA polymerase. The reaction was subjected to a 2-minute initial denaturation at 98°C, followed by 30 cycles of: 98°C for 10s, 68°C for 15s and 72°C for 10s, followed by a final extension step at 72°C of 2 minutes and cooling down to 10°C. The amplicons were purified using NucleoSpin Gel and PCR Clean-up kit (Macherey-Nagel) following the manufacturer’s instructions.

### In vitro production and purification of RNA

RNA was produced by in vitro transcription. In a volume of 500 µL, 2-3 µg purified DNA template was mixed with transcription buffer (40 mM HEPES pH 7.5, 20 mM MgCl2, 50 mM KCL, 2 mM Spermidine), 10 mM NTPs (2.5 mM each), 10 mM DTT, 0.4 U/µL RiboLock Inhibitor (Thermo Scientific) and in-house produced T7 RNA polymerase. The reaction was incubated at 37°C overnight and stopped by adding 2 Units of DNase I (NEB) and incubating at 37°C for 15 minutes. The reactions were centrifuged at 17,000 RCF for 10 min to pellet the precipitated pyrophosphate. The supernatant was loaded onto a Superose 6 size exclusion column (GE Healthcare) equilibrated with 40 mM HEPES pH 7.5, 50 mM KCL and 5 mM MgCl2.

### Fluorescence measurements

Excitation and emission spectra of the aptamer-fluorophore complexes were identified with spectral scan measurements on a CLARIOstar Plus multi-mode microplate reader (BMG LABTECH). All fluorescence measurements were performed at room temperature on sample volumes of 50 µl containing 100 nM RNA, 500 nM DFHBI-1T (Lucerna Technologies), 500 nM HBC620 (FR Biotechnology), 40 mM HEPES, 50 mM KCl and 5 mM MgCl2 using a CLARIOstar Plus multi-mode microplate reader (BMG LABTECH). Excitation of DFHBI-1T was performed at 460 nm and emission was recorded at 505 nm. Excitation of HBC620 was performed at 580 nm and emission was recorded at 620 nm. Fluorescence coming from the FRET was obtained by exciting at 460 nm and collecting at 620 nm.

### FRET output calculation

FRET was calculated using the following formula ^6^:

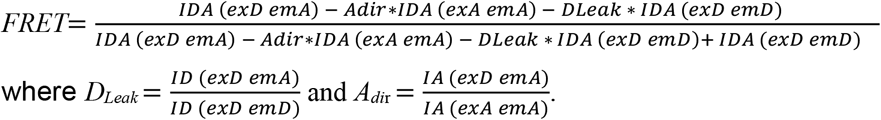

The excitation at DFHBI-1T or HBC620 wavelength is denoted with exD (460 nm) or exA (585 nm), respectively. The emission measured at DFHBI-1T or HBC620 wavelength is denoted with emD (505 nm), emA (620 nm), respectively. ID, IA and IDA refer to intensities measured in the presence of DFHBI-1, HBC620 and both fluorophores respectively.

### Cryo-EM sample preparation

Purified RNA (pre-incubated with fluorophores at a 1:5 molar ratio, or not) was spin concentrated to ∼2.5 mg/ml in Amicon centrifugal filters with molecular cuttoff weights of 10 kDa at 21 °C. Protochips AU-Flat 1.2/1.3 300 mesh grids were purchased from Jena Bioscience. Immediately prior to use the grids were glow discharged for 45 seconds with a current of 15 mA in a Pelca EasiGlow. A Leica GP2 was used for plunge-freezing, the sample chamber was kept at 15 degrees and 100% humidity. 3 μL of sample was applied to the gold foil and, after a delay of 4 seconds, blotted onto a double layer of Whatman number 1 filter paper for 6 seconds of total blot time followed by immediate plunging into liquid ethane (∼ - 184 °C).

### Cryo-EM data collection and single particle analysis

All data were acquired at 300 keV on a Titan Krios G3i (Thermo Fisher Scientific) equipped with a K3 camera (Gatan/Ametek) and energy filter operated in EFTEM mode using a slit width of 20 eV. The data were collected over a defocus range of -0.5 to -2 micrometers with a targeted dose of 60 e-/Å2. Automated data collection was performed with EPU and the data saved as gain normalized compressed tiff files with a pixel size of 0.645 Å/px.

All data were pre-processed using CS-Live to apply motion correction, CTF fitting and initial particle picking ^39^. The rest of the analysis was performed in cryoSPARC V3.31. For the ligand bound dataset templated particle picking using 50 templates generated from an ab initio reconstruction resulted in 729,630 particles, which were extracted with box size of 592 and Fourier cropped to 174 pixels. 3-class ab initio reconstruction using 30,000 particles resulted in 2 junk and 1 good class. Heterogeneous refinement was used to sort the particle stack into 1 good (reaching Nyquist) and 2 junk classes (see workflow in Supplementary Fig. 4). The 241,297 particles from the “good” class were used for 3D Variability Analysis (3DVA) solving for 3 orthogonal principal modes and a filter resolution of 7 Angstrom ^40^. These 241,297 particles were re-extracted with a box size of 592 and Fourier cropped to 296 pixels. The resultant 233,171 particles were used to start a 5-class ab initio reconstruction followed by heterogeneous refinement using the 5 ab initio reconstructions. The two best classes, totaling 150,204 particles, were combined and refined by homogeneous refinement followed by a local refinement using the mask from the homogeneous refinement job to attain the final particle alignments and reconstruction. Re-extracting the particles with a box size of 432, Fourier cropped to 216 pixels, and repeating the last two refinement steps improved both the map quality and GSFSC curve.

For the apo dataset a similar workflow was followed, resulting in 478,981 particle picks. A single round of 3D classification with 3-class ab initio reconstruction followed by heterogeneous refinement resulted in a refined particle stack of 51,278 particles. These particles were re-extracted with a box size of 432 and Fourier cropped to 216 pixels. Homogeneous refinement followed by local refinement with the mask from the previous homogeneous refinement job was used to attain the final particle alignments and reconstruction (see workflow in Supplementary Fig. 5).

The Local Resolution Estimation job in cryoSPARC was used to generate a local resolution mask that was applied to a locally filtered map in chimeraX.

### Model building

The core components of the RNA origami scaffold were generated using the ROAD software. The kissing loops were replaced with the kissing loop from helix 3 of another RNA origami (PDB: 7PTQ). The iSpinach aptamer (PDB: 5OB3) ^41^ was used as a starting template for Broccoli and the crystal structure of the Pepper aptamer bound to HBC620 (PDB: 7EOP) ^34^ was used as the starting template for Pepper. The components were manually placed into the cryo-EM volume in ChimeraX ^42-44^, the individual components were joined using the “make bond” command from the ISOLDE ^45^ add-on to ChimeraX. The resulting PDB file was re-numbered using the PDB-Tools pdb-reres program ^46^ and then the correctly numbered PDB file was sequence corrected in ChimeraX using the swapNA command. The model was then massaged into the cryo-EM volume using Molecular Dynamics Flexible Fitting (MDFF) with VMD using ISOLDE with a “temperature” of 0 degrees and a substantially reduced forcefield weight set ^45^. This model was then passed through real space refinement (RSR) in Phenix ^47-49^ using default parameters to optimize the backbone angles. As a final step the model was energy minimized with QRNAS ^50^ and iterated between Phenix RSR and QRNAS with positional restraints to allow the regions with bad clashes to be modified by QRNAS. Validation of the goodness of fit between model and map were performed using the Phenix validation tool ^49,51,52^.

### SHAPE analysis of the Pepper aptamer

For the investigation of the secondary structure of the Pepper aptamer, the construct with the F30 scaffold ^33^ fused to Pepper was selected. RNA was transcribed from a PCR amplified template containing three guanosyl residues at the 5’-end to facilitate transcription using T7 polymerase. For this, a long single stranded oligonucleotide (F30_Pepper_tplt) was designed as a template containing the full sequence and was amplified with two oligonucleotides (T7_Prom_Fwd and F30_Pepper_Rev) using Q5 High-fidelity DNA polymerase (NEB) according to the manufacturer’s instructions. 50 µL of the PCR reaction were used for a 500 µL in vitro transcription with T7 RNA polymerase (produced in-house). The transcription mix contain 200 mM Tris-HCl (pH=8), 40 mM DTT, 16 mM NTPs (4 mM each), 20 mM MgCl2, 2 mM spermidine, 10 µL of T7 polymerase and 20,000 units of RNase inhibitor (ThermoFischer). Transcription was run overnight at 37°C and RNA was purified on a 6% denaturing PAGE.

After purification, the RNA concentration was determined using DS 11 Series spectrophotometer (Denovix). For each reaction, 10 pmol of RNA was used. In total, 5 different reactions were setup. A control reaction (mock) that did not undergo any SHAPE treatment, a sample without HBC dye (neg) and three samples containing one of the fluorophores (HBC 485, 497 or 620), respectively. All RNAs were mixed in a buffer containing 40 mM HEPES pH 7, 50 mM KCL, 5 mM MgCl2 and incubated at 65°C for 5 min, followed by a 5 min incubation at room temperature. 4 µL of dye at 50 µM was mixed in their corresponding tubes (485, 497 and 620) and 4 µL of DMSO anhydrous was added to the mock and neg samples. Samples were incubated 5 min at room temperature before each being transferred into an Eppendorf tube containing either DMSO (mock) or 5 µL of 2 M Benzoyl Cyanide. As the chemical modification is done after 1 s ^53^, the samples were ethanol precipitated and resuspended into 9 µL of MQ water.

For the primer extension, a 2X master mix was prepared containing 2.5X SSII buffer, 500 µM dNTPs, 1.5 µM Alex647 modified reverse primer (Rev_Shape_Alexa647) and 20 mM DTT. 10 µL of this mix was added to the 9 µL of RNA. For the sequencing lanes of the gel, 4 samples containing 10 pmol of RNA were prepared, containing each a different ddNTP at 500 µM. All the samples were heated up at 65°C for 5 min, then 5 min at 35°C followed by 5 min at 25°C. 1 µL of SSII (ThermoFischer) was added to each tube. The samples were incubated 1 min at 45°C, 20 min at 52°C and 5 min at 65°C before being kept at 4°C. Once the cDNA synthesis was over, the remaining RNA was degraded by adding 1 µL of 4 M NaOH and incubated 5 min at 95°C. All samples were ethanol precipitated and resuspended in 15 µL of loading buffer (97% formamide and 20 mM EDTA).

For the observation of the result, a 12% PAGE (40 cm x 20 cm x 0.07 cm) was cast. Prior loading, all samples were heated up at 95°C for 3 min before being snapped cool on ice for 3 min. After 15 min of running at 1000 V, 3 µL of the samples (mock, neg, 485, 497 and 620) and 1.5 µL of the sequencing samples were loaded on the gel. After 5h of migration, the gel was scanned on a Typhoon FLA-9500.

Reactivity was calculated by measuring the peak intensity of each band using the ImageJ software. Each sample was normalized based on their signal intensity of the C28. This nucleotide is located in the P1 apical tetraloop, far from the binding region and is assumed to have no reactivity variation upon target binding due to the inherent stability of a UUGC tetraloop.

## DATA AVAILABILITY

The atomic coordinates for the 1,2-B12P12 RNA origami scaffolds in the bound and apo states have been deposited in the PDB (https://www.rcsb.org/) under the PDB ID 7ZJ4 and 7ZJ5, respectively. The volumes from the final refinements of our cryo-EM SPA datasets have been deposited to the ePDB under accession codes EMDB-14740 and EMDB-17471. Other data are available from the corresponding author upon request.

## FUNDING

N. S. V. received funding from the European Union’s Horizon 2020 Research and Innovation Program under the Marie Sklodowska-Curie grant agreement n° 765703. E. K. S. M. was supported by the Independent Research Fund Denmark under the Research Project 1 grant (9040-00425B) and the Canadian Natural Sciences and Engineering Research Council (532417). Computational resources for the project were in part supported by the Carlsberg Foundation Research Infrastructure grant (CF20-0635). B. K. H. was supported by a PhD scholarship from Innovation Foundation Denmark. E.S.A. acknowledges support by a European Research Council (ERC) Consolidator grant (683305) and Novo Nordisk Foundation Ascending Investigator grant (0060694) supporting A.B.

## AKNOWLEDGEMENTS

We thank Mette Jepsen for insightful discussions and Rita Rosendahl and Claus Bus for technical assistance.

## CONTRIBUTION

N. S. V., B. K. H., E. K. S. M. and E. S. A. conceptualized the project and designed the experiments. N. S. V. and B. K. H. designed the RNAs and performed the FRET experiments. E. K. S. M. performed the cryo-EM characterization and analysis. A. B. performed the SHAPE probing experiments. N. S. V., E. K. S. M. and E. S. A. wrote the manuscript.

## ETHICS DECLARATION

The authors declare no competing interests.

## Notes

### Competing Interest Statement

The authors have declared no competing interest.

