## Supporting Information for "RNA origami scaffolds as a cryo-EM tool for investigating aptamer-ligand binding of a Broccoli-Pepper FRET pair"

**Supplementary Table 1: RNA designs and sequences.**

| 1,2-B12P12 |
|---|
| 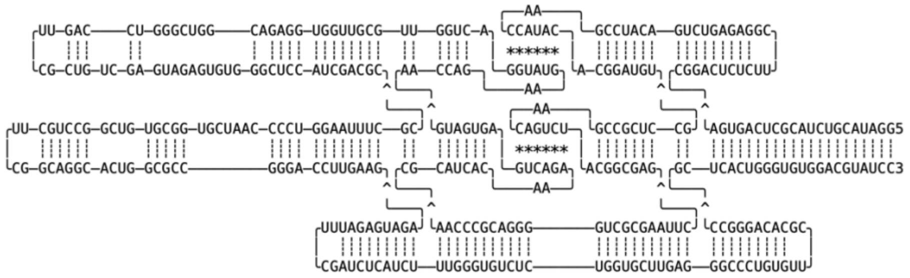 |
| GGAUACGUCUACGCUCAGUGACGGACUCUCUUCGGAGAGUCUGACAUCCGAACCAUACACGGAUGUGCCUCGCCGAACAGUCUACGGCGAGCUUAAG<br>CGCUGGGGACGCCCAACGCAUCACAAAGACUGAGUGAUGAACCAGAAGUAUGGACUGGUUGCGUUGGUGGAGACGGUCGGGUCCAGUUCGCUGUCGA<br>GUAGAGUGUGGGCUCCAUCGACGCCGCUUUAAGGUCCCCAAUCGUGGCGUGUCGGCCUGCUUCGGCAGGCACUGGCGCCGGGACCUUGAAGAGAUGA<br>GAUUUCGAUCUCAUCUUUGGGUGUCUCUGGUGCUUGAGGGCCCUGUGUUCGCACAGGGCCGCUCACUGGGUGUGGACGUAUCC |

| 1,3-B12P12 |
|---|
| 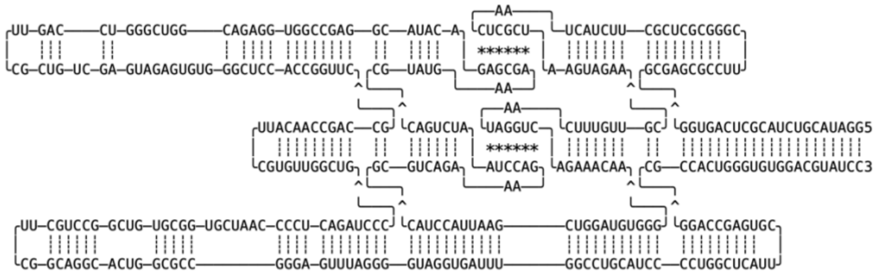 |
| GGAUACGUCUACGCUCAGUGGGCGAGCGCCUUCGGGCGCUCGCUUCUACUAACUCGCUAAGUAGAACGUUGUUUCAAUAGGUCAGAAACAAGGGUGU<br>AGGUCGAAUUACCUACGCGUCAGAAAGACCUAAUCUGACCGUAUGAAAGCGAGACAUACGGAGCCGGUGGAGACGGUCGGGUCCAGUUCGCUGUCGA<br>GUAGAGUGUGGGCUCCACCGGUUCGCCAGCCAACAUUCGUGUUGGCUGCCCUAGACUCCCCAAUCGUGGCGUGUCGGCCUGCUUCGGCAGGCACUGG<br>CGCCGGGAGUUUAGGGGUAGGUGAUUUGGCCUGCAUCCCCUGGCUCAUUCGUGAGCCAGGCGCCACUGGGUGUGGACGUAUCC |

| 1,2-B12P10 |
|---|
| 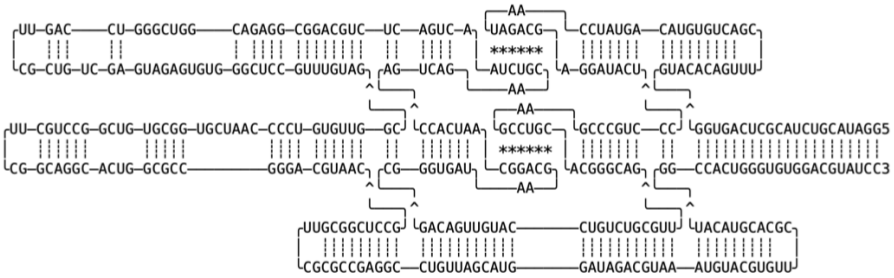 |
| GGAUACGUCUACGCUCAGUGGGUACACAGUUUCGACUGUGUACAGUAUCCAAUAGACGAGGAUACUCCCUGCCCGAAGCCUGCACGGGCAGUUGCGU<br>CUGUCCAUGUUGACAGCGGGUGAUAAGCAGGCAAUCACCAGUCAGAACGUCUAACUGACUCUGCAGGCGGAGACGGUCGGGUCCAGUUCGCUGUCGA<br>GUAGAGUGUGGGCUCCGUUUGUAGCGGUUGUGUCCCCAAUCGUGGCGUGUCGGCCUGCUUCGGCAGGCACUGGCGCCGGGACGUAACGCCUCGGCGU<br>UCGCGCCGAGGCCUGUUAGCAUGGAUAGACGUAAAUGUACGUGUUCGCACGUACAUGGCCACUGGGUGUGGACGUAUCC |

**Supplementary Table 1: RNA designs and sequences (continued).**

| 1,2-B12P14 |
| --- |
| 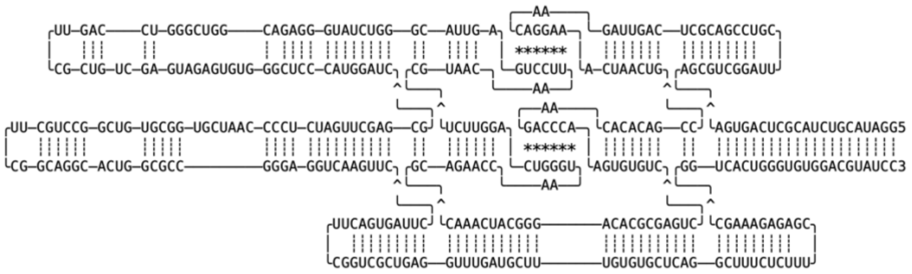 |
| GGAUACGUCUACGCUCAGUGAAGCGUCGGAUUCGUCCGACGCUCAGUUAGAACAGGAAACUAACUGCCGACACACAAGACCCAAGUGUGUCCUGAGC GCACAGGGCAUCAAACGCAGAACCAAUGGGUCAGGUUCUCGUAACAAUUCCUGAGUUACGGGUCUAUGGGAGACGGUCGGGUCCAGUUCGCUGUCGA GUAGAGUGUGGGCUCCCAUGGAUCGCGAGCUUGAUCUCCCCAAUCGUGGCGUGUCGGCCUGCUUCGGCAGGCACUGGCGCCGGGAGGUCAAGUUCCU UAGUGACUUCGGUCGCUGAGGUUUGAUGCUUUGUGUGCUCAGGCUUUCUCUUUCGAGAGAAAGCGGUCACUGGGUGUGGACGUAUCC |

| 1,3-B12P(-34) |
| --- |
| 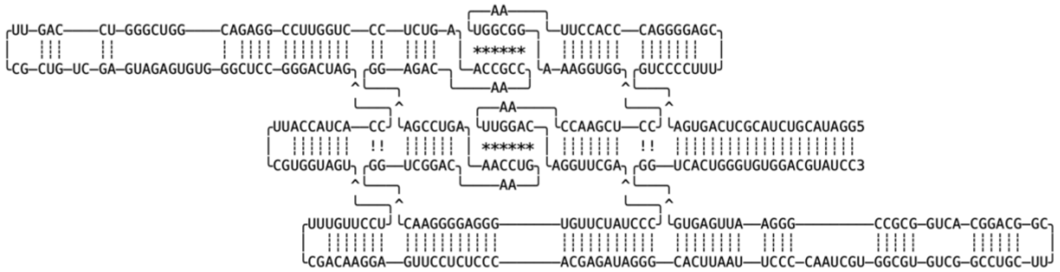 |
| GGAUACGUCUACGCUCAGUGAGUCCCCUUUCGAGGGGACCCACCUUAAUGGCGGAAAGGUGGCCUCGAACCAAUUGGACAGGUUCGACCCUAUCUUG UGGGAGGGGAACGGUCGGACAAGUCCAAAGUCCGAGGAGACAACCGCCAAGUCUCCCUGGUUCCGGAGACGGUCGGGUCCAGUUCGCUGUCGAGUAG AGUGUGGGCUCCGGGACUAGCCACUACCAUUCGUGGUAGUUCCUUGUUUCGACAAGGAGUUCCUCUCCCACGAGAUAGGGCACUUAAUUCCCCAAUC GUGGCGUGUCGGCCUGCUUCGGCAGGCACUGGCGCCGGGAAUUGAGUGGGUCACUGGGUGUGGACGUAUCC |

| F30-Pepper |
| --- |
| 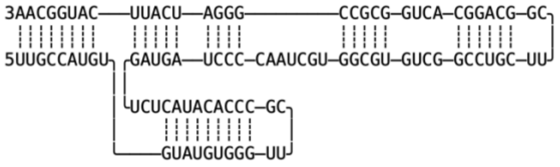 |
| UUGCCAUGUGUAUGUGGGUUCGCCCACAUACUCUGAUGAUCCCCAAUCGUGGCGUGUCGGCCUGCUUCGGCAGGCACUGGCGCCGGGAUCAUUCAUG GCAA |

**Supplementary Table 2. Fluorescence intensities from spectrofluorometric measurements and FRET calculations.**

(A) Fluorescence intensities measured at excitation and emission maxima. The excitation of DFHBI-1T or HBC620 is denoted with $ex_D$ (460 nm) or $ex_A$ (585 nm), respectively. The emission measured of DFHBI-1T or HBC620 is denoted with $em_D$ (505 nm), $em_A$ (620 nm), respectively. $I_D$, $I_A$ and $I_{DA}$ refer to intensities measured in the presence of DFHBI-1T, HBC620 and both fluorophores, respectively. Data corresponds to mean value ± standard error from three pipetting replicates. (B) Calculations of donor leak ($D_{Leak}$), direct acceptor excitation ($A_{dir}$) and FRET output. Data corresponds to mean value ± standard error from three technical replicates.

**A**

| | $I_D$ ($ex_D$ $em_D$) | $I_D$ ($ex_D$ $em_A$) | $I_A$ ($ex_D$ $em_A$) | $I_A$ ($ex_A$ $em_A$) | $I_{DA}$ ($ex_D$ $em_D$) | $I_{DA}$ ($ex_D$ $em_A$) | $I_{DA}$ ($ex_A$ $em_A$) |
|---|---|---|---|---|---|---|---|
| **1, 2-B12P12** | 40230.7 ± 847 | 281.3 ± 4.7 | 1404.7 ± 22.2 | 28540 ± 781.2 | 9459 ± 225.1 | 7042 ± 755.9 | 27876.7 ± 677.3 |
| **1, 3-B12P12** | 38834.3 ± 587.4 | 257.3 ± 5.5 | 1171.6 ± 26 | 25582.7 ± 326.5 | 22492 ± 331.5 | 3771 ± 63.6 | 25386.3 ± 400.9 |
| **1, 2-B12P10** | 36290.7 ± 91 | 244.7 ± 15.1 | 1187 ± 41.2 | 24147 ± 679.2 | 8233.3 ± 110.7 | 5541 ± 88.4 | 23485.3 ± 501.2 |
| **1, 2-B12P14** | 38828.3 ± 779.9 | 291 ± 22.1 | 1310 ± 12.8 | 25633.3 ± 602.9 | 11236.7 ± 206.3 | 5932.3 ± 104.7 | 25373 ± 460.4 |
| **1,3-B12-P(-34)** | 37365 ± 1168.2 | 289 ± 15.1 | 101.7 ± 66.8 | 24469 ± 767.1 | 32691.7 ± 780.6 | 1332.7 ± 45.3 | 24602 ± 552.1 |

**B**

| | $D_{Leak}$ | $A_{dir}$ | FRET output |
|---|---|---|---|
| **1, 2-B12P12** | 0.7 ± 0.01 % | 4.93 ± 0.08 % | 0.372 ± 0.004 |
| **1, 3-B12P12** | 0.66 ± 0.01 % | 4.58 ± 0.05 % | 0.099 ± 0.0002 |
| **1, 2-B12P10** | 0.67 ± 0.04 % | 4.91 ± 0.07 % | 0.345 ± 0.0004 |
| **1, 2-B12P14** | 0.75 ± 0.05 % | 5.12 ± 0.09 % | 0.288 ± 0.001 |
| **1,3-B12-P(-34)** | 0.77 ± 0.04 % | 4.08 ± 0.16 % | 0.002 ± 0.002 |

**Supplementary Table 3. Cryo-EM data collection, refinement and validation statistics.**

| | #1 Apta_FRET_Bound (EMDB-14740) (PDB 7ZJ4) | #2 Apta_FRET_Apo (EMDB-17471) (PDB 7ZJ5) |
|---|---|---|
| **Data collection and processing** | | |
| Magnification | 130000 | 130000 |
| Voltage (kV) | 300 | 300 |
| Electron exposure (e–/Å$^2$) | 60 | 60 |
| Defocus range (μm) | -0.7 to -2 | -0.7 to -2 |
| Pixel size (Å) | 0.647 | 0.647 |
| Symmetry imposed | none | none |
| Initial particle images (no.) | 729630 | 478981 |
| Final particle images (no.) | 150204 | 51278 |
| Map resolution (Å) | 4.43 | 4.55 |
| FSC threshold | | |
| Map resolution range (Å) | 3.77-8.89 | 4.3-10.6 |
| **Refinement** | | |
| Initial model used (PDB code) | 7PTQ - 7EOP | 7PTQ - 7EOP |
| Model resolution (Å) | 4.4 (0.143) | 4.5 (1.43) |
| FSC threshold | | |
| Map sharpening *B* factor (Å$^2$) | 174 | 158 |
| Model composition | | |
| Non-hydrogen atoms | 8026 | 7984 |
| Nucleotide residues | 374 | 374 |
| Ligands | 3 | 1 |
| *B* factors (Å$^2$) | (mean) | (mean) |
| Nucleotide | 234 | 423 |
| Ligand | 461 | 1012 |
| R.m.s. deviations | | |
| Bond lengths (Å) | 0.004 (0) | 0.002 (0) |
| Bond angles (°) | 0.837 (0) | 0.652 (0) |
| Validation | | |
| MolProbity score | 1.98 | 1.82 |
| Clashscore | 1.16 | 0.50 |

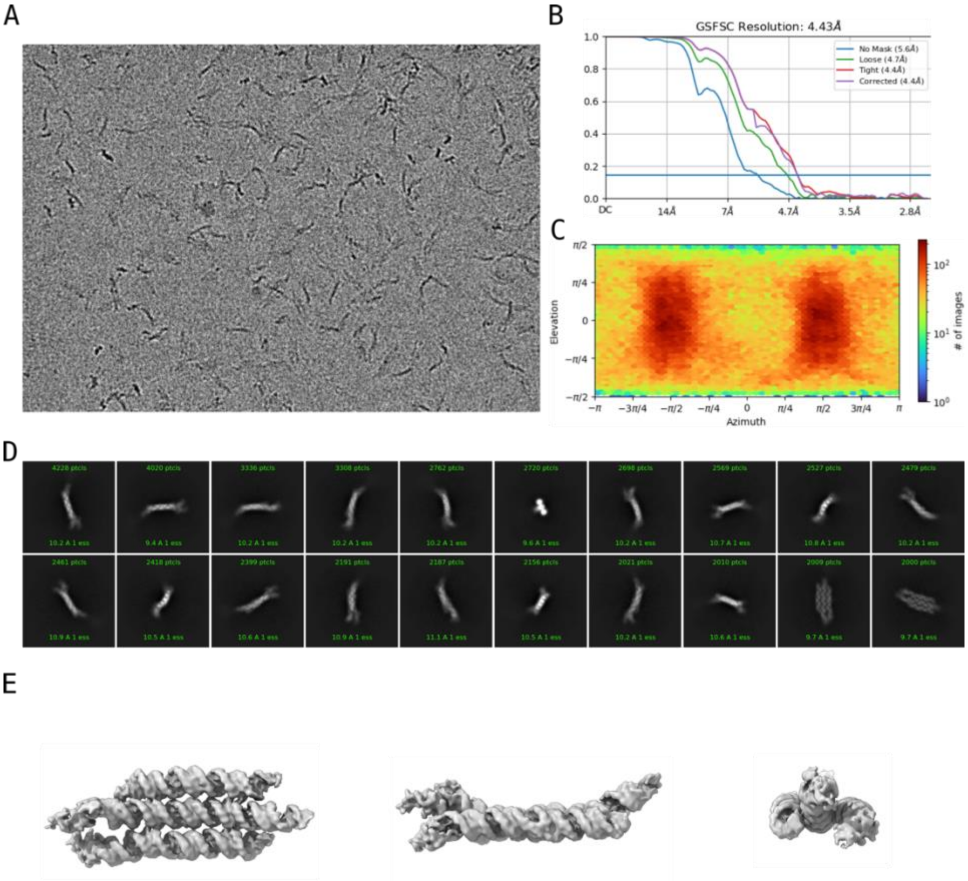

**Supplementary Fig. 1. Cryo-EM data and reconstruction of ligand bound 1,2-B12P12.** (A) Example cryo-EM micrograph from the ligand bound 1,2-B12P12 dataset. (B) Gold-Standard Fourier Shell Correlation for the ligand bound 1,2-B12P12 reconstruction. (C) Angular distribution of particles used in final reconstruction. (D) 2D classes from the final particle stack of the ligand bound 1,2-B12P12 dataset. (E) Three alternate views of the ligand bound 1,2-B12P12 reconstruction.

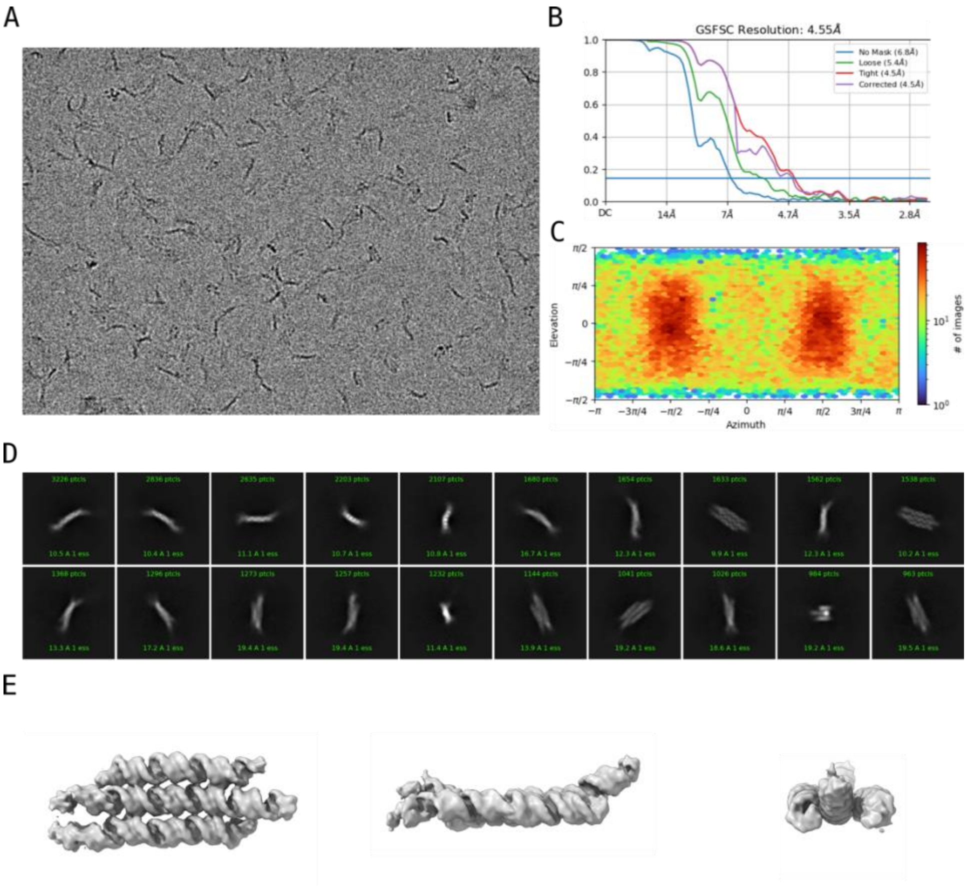

**Supplementary Fig. 2. Cryo-EM data and reconstruction of the Apo 1,2-B12P12.**
(A) Example cryo-EM micrograph from the Apo 1,2-B12P12 dataset. (B) Gold-Standard Fourier Shell Correlation for the Apo 1,2-B12P12 reconstruction. (C) Angular distribution of particles used in final reconstruction. (D) 2D classes from the final particle stack of the Apo 1,2-B12P12 dataset. (E) Three alternate views of the Apo 1,2-B12P12 reconstruction.

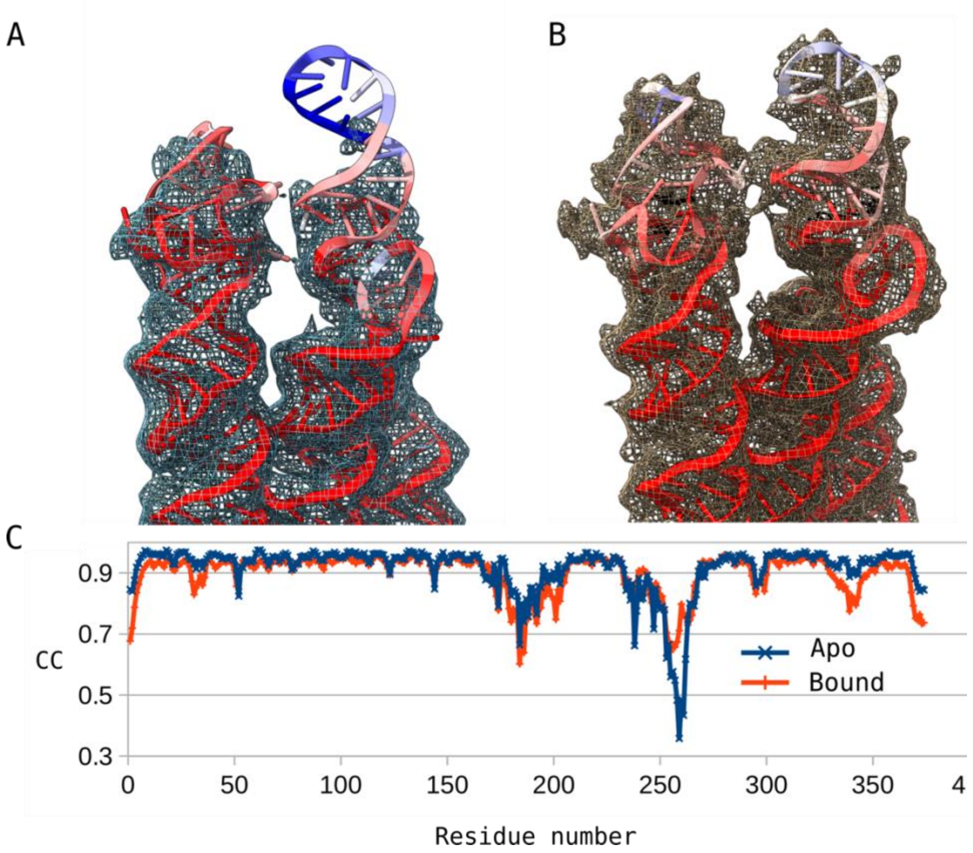

**Supplementary Fig. 3. Goodness of fit for the aptamers modelled into the Apo and Bound reconstructions.**

Broccoli and Pepper models colored by per-residue cross correlation coefficients (CC) for the Apo (A) and Bound (B) reconstructions. Coloring of the models is on a scale from 0.3-0.9 from blue to red, reconstructions are shown as a mesh surface with threshold level set to 0.091. (C) Per-residue CC is plotted vs residue number for the Apo and Bound models.

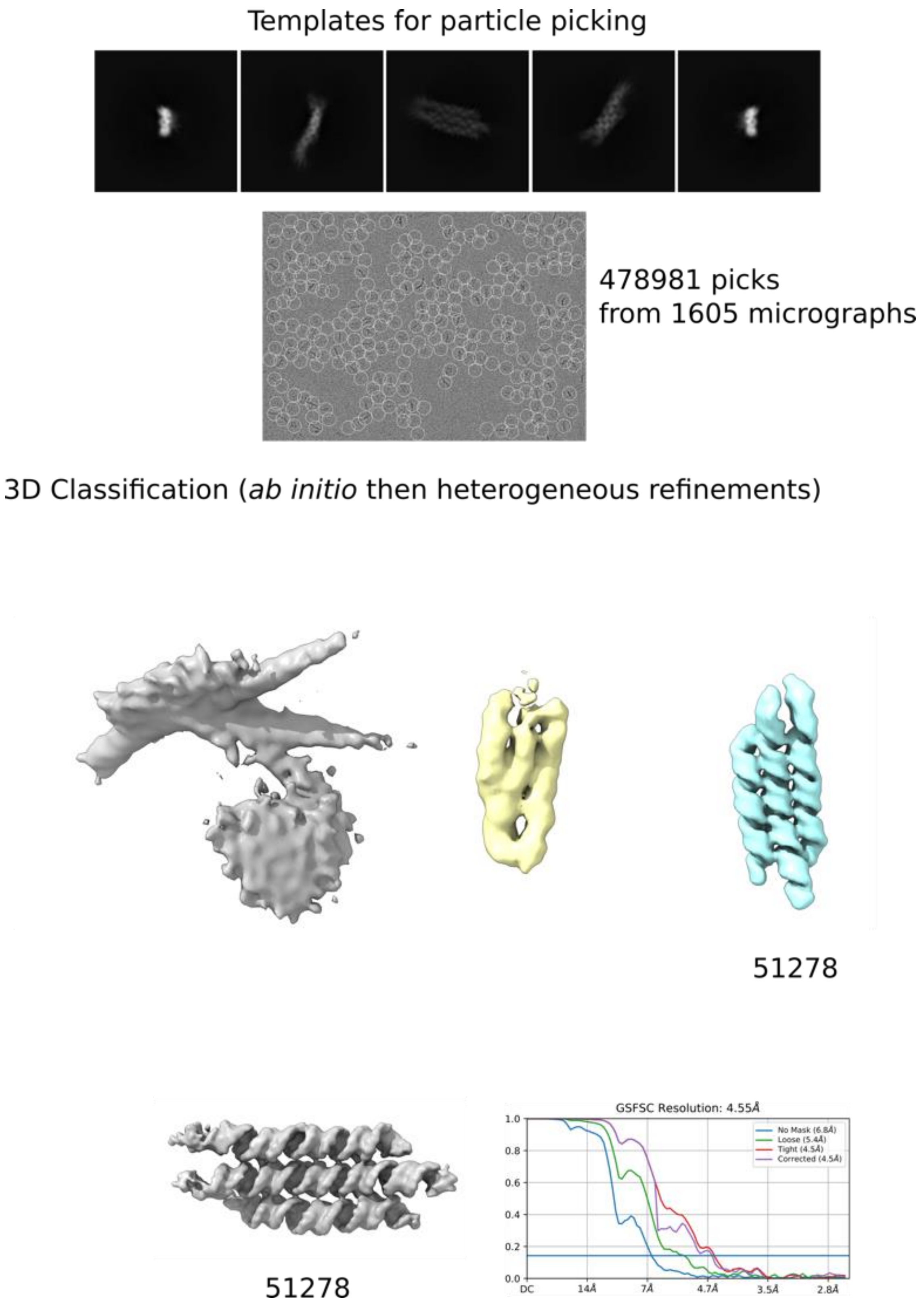

**Supplementary Fig. 4. Single particle analysis workflow for the Apo 1,2-B12P12 RNA.** 2D templates generated from the ab initio reconstruction obtained during a CS-Live session were used to re-pick particles from the motion and CTF corrected micrographs, followed by classification in 3D to attain the final particle stack.

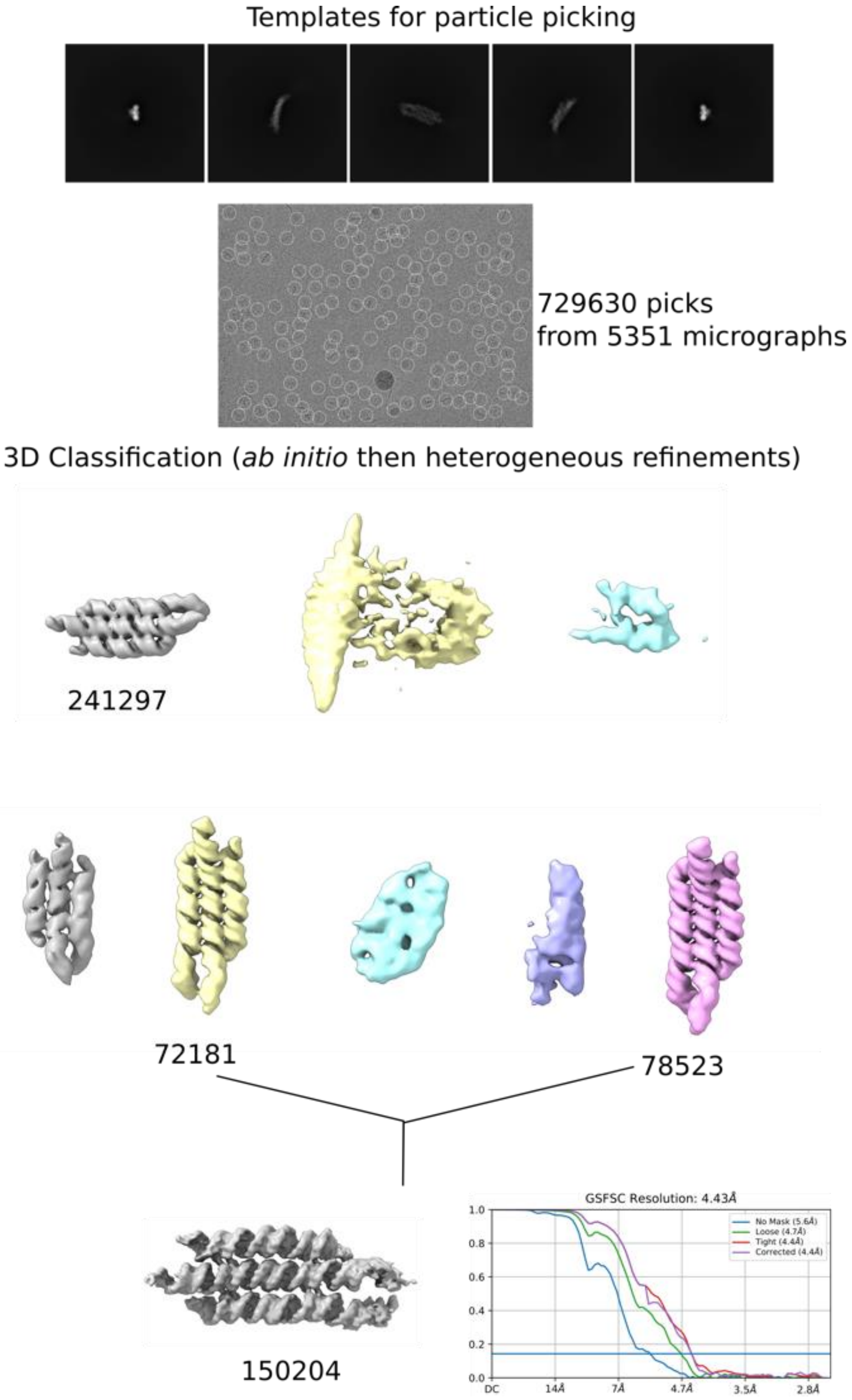

**Supplementary Fig. 5. Single particle analysis workflow for the ligand bound 1,2-B12P12 RNA.**
2D templates generated from the ab initio reconstruction obtained during a CS-Live session were used to re-pick particles from the motion and CTF corrected micrographs, followed by classification in 3D to attain the final particle stack.

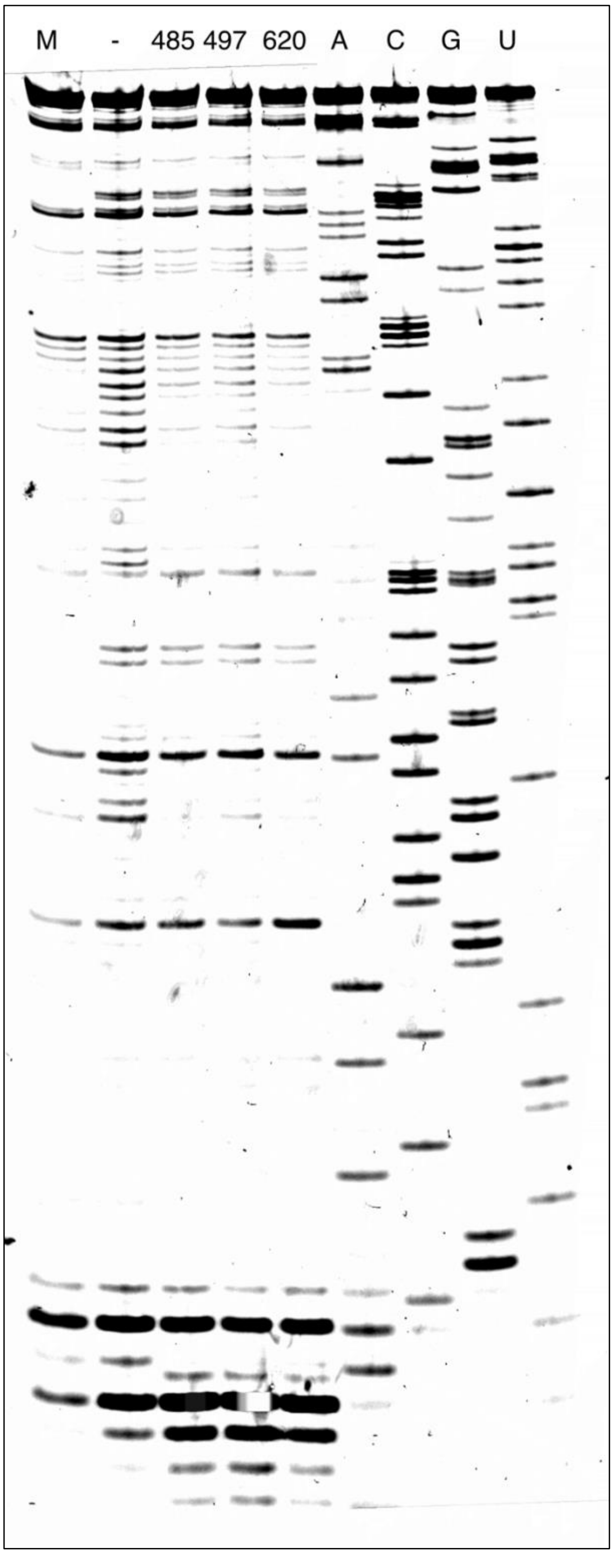

102

103 **Supplementary Fig. 6. Full image of the SHAPE probing PAGE gel.**

104
